# Human VMPFC encodes early signatures of confidence in perceptual decisions

**DOI:** 10.1101/224337

**Authors:** Sabina Gherman, Marios G. Philiastides

## Abstract

Choice confidence, an individual’s internal estimate of judgment accuracy, plays a critical role in adaptive behaviour. Despite its importance, the early (decisional) stages of confidence processing remain underexplored. Here, we recorded simultaneous EEG/fMRI while participants performed a direction discrimination task and rated their confidence on each trial. Using multivariate single-trial discriminant analysis of the EEG, we identified a stimulus- and accuracy-independent component encoding confidence, appearing prior to subjects’ choice and explicit confidence report. The trial-to-trial variability in this EEG-derived confidence signal was uniquely associated with fMRI responses in the ventromedial prefrontal cortex (VMPFC), a region not typically associated with confidence for perceptual decisions. Furthermore, we showed that the VMPFC was functionally coupled with regions of the prefrontal cortex that support neural representations of confidence during explicit metacognitive report. Our results suggest that the VMPFC encodes an early confidence readout, preceding and potentially informing metacognitive evaluation and learning, by acting as an implicit value/reward signal.

## Introduction

Our everyday lives involve frequent situations where we must make judgments based on noisy or incomplete sensory information – for example deciding whether crossing the street on a foggy morning, in poor visibility, is safe. Being able to rely on an internal estimate of whether our perceptual judgments are accurate is fundamental to adaptive behaviour and accordingly, recent years have seen a growing interest in understanding the neural basis of confidence judgments.

Within the perceptual decision making field, one line of research has focused specifically on identifying neural correlates of confidence during metacognitive evaluation (i.e., while subjects actively judge their performance following a choice), and demonstrated the functional involvement of the anterior prefrontal cortex (Fleming et al., 2012; Hilgenstock et al., 2014). Concurrently, psychophysiological work in humans and non-human primates using time-resolved measurements have shown that confidence encoding can also be observed at earlier stages, and as early as the decision process itself (Kiani and Shadlen, 2009; Zizlsperger et al., 2014; Gherman and Philiastides, 2015).

Correspondingly, recent fMRI studies have reported confidence-related signals nearer the time of decision (e.g., during perceptual stimulation) in regions such as the striatum (Hebart et al., 2016), dorsomedial prefrontal cortex (Heereman et al., 2015), cingulate and insular cortices (Paul et al., 2015), and other areas of the prefrontal, parietal, and occipital cortices (Heereman et al., 2015; Paul et al., 2015). Interestingly, confidence-related processing has also been reported in the ventromedial prefrontal cortex (VMPFC) during value-based and a range of ratings tasks (De Martino et al., 2013; Lebreton et al., 2015), however the extent to which this region is additionally involved in perceptual judgments relying on temporal integration of sensory evidence remains unclear.

Importantly, research investigating the neural correlates of decision confidence has thus far relied – nearly exclusively – on correlations with behavioural measures, the most common of these being the subjective ratings given by participants after the decision (see (Grimaldi et al., 2015) for a review). However, theoretical and empirical work suggests that post-decisional metacognitive reports may be affected by processes occurring after termination of the initial decision (Resulaj et al., 2009; Pleskac and Busemeyer, 2010; Fleming et al., 2015; Moran et al., 2015; Murphy et al., 2015; Yu et al., 2015; Navajas et al., 2016; van den Berg et al., 2016; Fleming and Daw, 2017), such as integration of existing information, processing of novel information arriving post-decisionally, or decay (Moran et al., 2015), and may consequently be only partly reflective of early confidence-related states.

Here we aim to derive a more faithful representation of these early confidence signals using EEG, and exploit the trial-by-trial variability in these signals to build parametric EEG-informed fMRI predictors, thus aiming to provide a more complete spatiotemporal account of decision confidence. We hypothesise that using an electrophysiologically-derived (i.e. endogenous) representation of confidence to detect associated fMRI responses would provide not only a more temporally precise, but also a more accurate spatial representation of confidence around the time of decision.

To test this hypothesis, we collected simultaneous EEG/fMRI data while participants performed a random-dot direction discrimination task and rated their confidence on each trial. Using a multivariate single-trial classifier to discriminate between High vs. Low confidence trials in the EEG data, we extracted an early, stimulus- and accuracy-independent discriminant component appearing prior to participants’ behavioural response. We then regressed the resultant single-trial component amplitudes against the fMRI signal and identified a positive correlation with this early confidence signal in a region of the VMPFC that has not been previously linked to perceptual decisions. Crucially, activation of this region was unique to our EEG-informed fMRI predictor (i.e., additional to those detected with a conventional fMRI regressor, which relied solely on participants’ post-decisional confidence reports). Furthermore, a functional connectivity analysis revealed a link between the activation in the VMPFC and regions of the prefrontal cortex found to hold neural representations of confidence during explicit metacognitive report.

## Results

### Behaviour

Subjects (N=24) performed a speeded perceptual discrimination task whereby they were asked to judge the motion direction of random dot kinematograms (left vs. right), and rate their confidence in each choice on a 9-point scale (Fig. 1A). Stimulus difficulty (i.e., motion coherence) was held constant across all trials, at individually determined psychophysical thresholds. We found that on average, subjects indicated their direction decision 994 ms (SD = 172 ms) after stimulus onset and performed correctly on 75% (SD = 5.2%) of the trials. In providing behavioural confidence reports, subjects tended to employ the entire rating scale, showing that subjective confidence varied from trial-to-trial despite perceptual evidence remaining constant throughout the task. As a general measure of validity of subjects’ confidence reports, we first examined the relationship with behavioural task performance. Specifically, confidence is largely known to scale positively with decision accuracy and negatively with response time (Vickers and Packer, 1982; Baranski and Petrusic, 1998) (though this relationship is not perfect, and is subject to individual differences, e.g., (Baranski and Petrusic, 1994; Fleming et al., 2010; Fleming and Dolan, 2012). As expected, we found a positive correlation with accuracy (subject-averaged R = .30; one-sample t-test, t(23) = 13.9, p < .001) (Fig. 1B), and a negative correlation with response time (subject-averaged R = -.27; one-sample t-test, t(23) = -7.8, p < .001) (Fig. 1C). Thus, subjects’ confidence ratings were generally reflective of their performance on the perceptual decision task.

**Figure 1.**
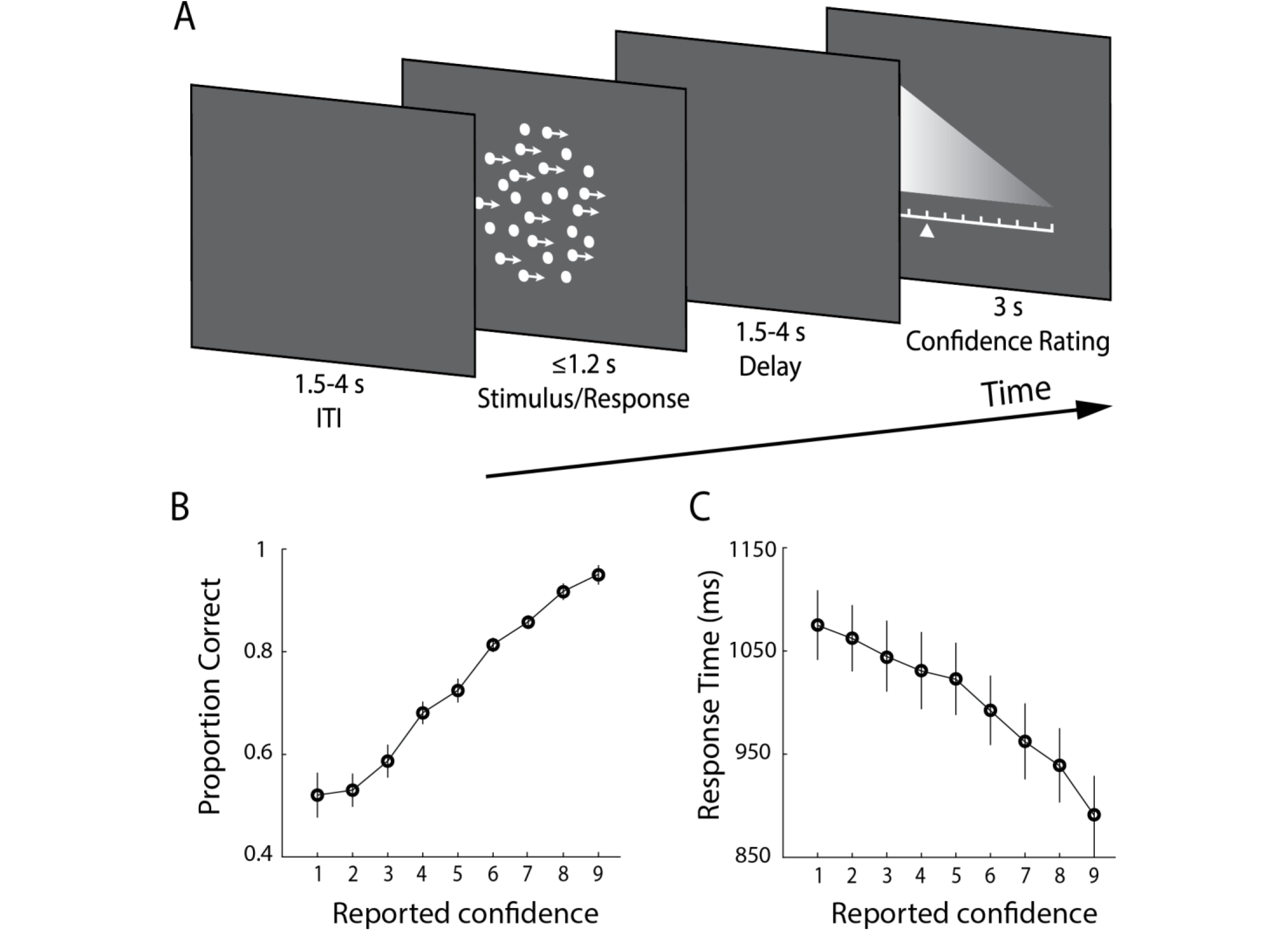
Experimental design and behavioural performance. A, Schematic representation of the behavioral paradigm. Subjects made speeded left vs. right motion discriminations of random dot kinematograms calibrated to each individual’s perceptual threshold. Stimulus difficulty (i.e., motion coherence) and was held constant across trials. Stimuli were presented for up to 1.2 s, or until a behavioural response was made. After each direction decision, subjects rated their confidence on a 9-point scale (3 s). The response mapping for high vs. low confidence ratings alternated randomly across trials to control for motor preparation effects, and was indicated by the horizontal position of the scale, with the tall end representing high confidence. All behavioural responses were made on a button box, using the right hand. B, Mean proportion of correct direction choices as a function of reported confidence. C, Mean response time as a function of reported confidence. Error bars in B and C represent the standard errors across subjects.

Next, we asked whether subjects’ confidence reports could be explained by local fluctuations in attention. To address this possibility, we performed a serial autocorrelation regression analysis on a single subject basis, which predicted confidence ratings on the current trial from ratings given on the immediately preceding five trials. On average, this model accounted for only a minimal fraction of the variance in confidence ratings (subject-averaged R^2^ = .07). Finally, we sought to rule out the possibility that trial-to-trial variability in confidence could be explained by potential subtle differences in low-level physical properties of the stimulus that may go beyond motion coherence (e.g., location and/or timing of individual dots). To this end, we compared subjects’ confidence reports on the two experimental blocks which contained an identical set of stimuli, and found no significant correlation between these (R = 0.02, p = 0.44). Taken together, these results support the hypothesis that subjects’ reports reflected internal fluctuations in their sense of confidence, which are largely unaccounted for by external factors.

### EEG-derived measure of confidence

To identify confidence-related signals in the EEG data, we first separated trials into three confidence groups (Low, Medium, and High) on the basis of subjects’ confidence ratings. We then conducted a single-trial multivariate classifier analysis (Parra et al., 2005; Sajda et al., 2009) on the stimulus-locked EEG data to estimate linear spatial weightings of the EEG sensors discriminating between Low vs. High confidence trials (see Materials and Methods). Applying the estimated electrode weights to single-trial data produced a measurement of the discriminating component amplitudes (henceforth *y*), which represent the distance of individual trials from the discriminating hyperplane, which we treat as a surrogate for the neural confidence of the decision. Note that separating trials in this manner only served to increase the precision of the discrimination process, i.e., estimate the electrode contribution patterns that optimally captured confidence. Data from all trials, including those not originally used in the discrimination analysis, were subsequently subjected through these spatial filters, resulting in discriminant component amplitudes that represent graded (individual trial) measures of internal confidence. To quantify the discriminator’s performance over time we used the area under a receiver operating characteristic curve (i.e. *Az* value) with a leave-one-out trial cross validation approach.

**Figure 2.**
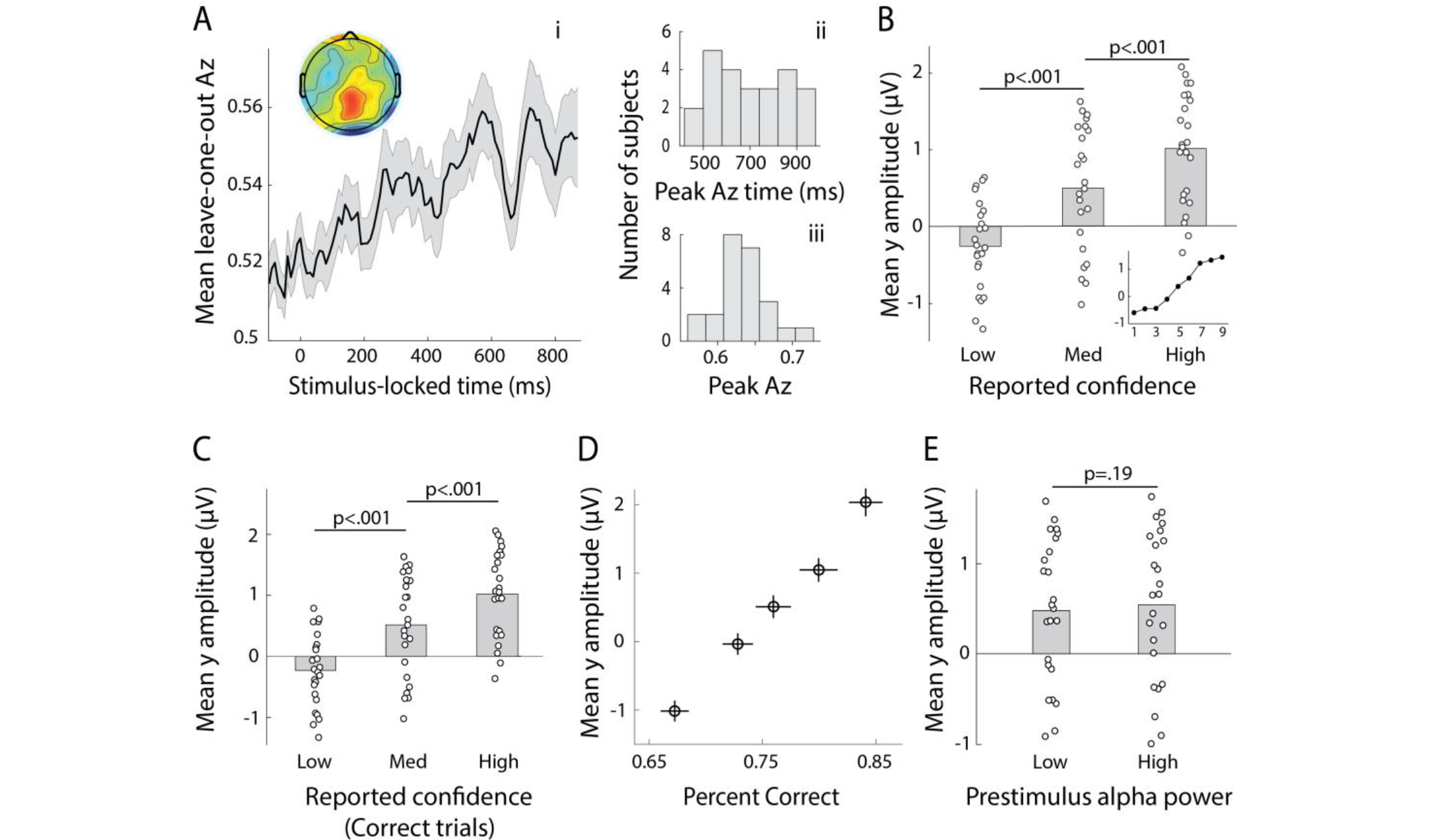
Neural representation of confidence in the EEG. **A**, Classifier performance (Az) during High-vs. Low-confidence discrimination for stimulus-locked single-trial data, **i**. Mean confidence discrimination performance as a function of time (shaded area represents standard errors across subjects). Inset shows average (normalised) topography associated with the discriminating component at subject-specific times of peak confidence discrimination, **ii**. Distribution of peak confidence discrimination times across subjects. In selecting these, we considered only the discrimination period ending on average at least 100 ms (across-subject mean 271±162 ms) prior to the subjects’ mean response times, to minimise potential confounds with activity related to motor execution (due to a sudden increase in corticospinal excitability in this period (Chen et al., 1998), iii. Distribution of Az values at the time of peak confidence discrimination across subjects. **B**, Mean amplitude of the confidence discriminant component as a function of confidence group (Low, Medium, High; grey bars). As expected, component amplitudes for the Medium confidence trials (i.e., trials which were independent from those used to perform the discrimination analysis) are situated between the Low and High confidence trials. The mean component amplitudes for individual confidence ratings (weighted by each subjects’ trial count per rating) are also shown (inset). **C**, Mean amplitude of the confidence discriminant component as a function of reported confidence, for correct trials only. The same pattern as in **B** is observed. **D**, Trial-by-trial confidence discriminant component amplitudes were positively correlated with accuracy. To visualise this relationship, single-trial component amplitudes were grouped into five bins. Error bars represent the standard error of the mean across subjects. **E**, Mean amplitudes of the confidence discriminant component did not differ significantly between trials associated with High vs. Low prestimulus oscillatory power in the alpha band. White dots in **B**, **C**, and **E** represent individual subject means.

We found that discrimination performance (Az) between the two confidence trial groups peaked, on average, 708 ms after stimulus onset (SD = 162ms, Fig. 2A). To visualise the spatial extent of this confidence component, we computed a forward model of the discriminating activity (Materials and methods), which can be represented as a scalp map (Fig. 2A). Importantly, both the temporal profile and electrode distribution of confidence-related discriminating activity appeared consistent with our previous work (Gherman and Philiastides, 2015) where we used stand-alone EEG to identify time-resolved signatures of confidence during a face vs. car task. Together these observations are an indication that the temporal dynamics of decision confidence can be reliably captured using EEG data acquired inside the MR scanner, and that these early confidence-related signals may generalise across tasks.

To provide additional support linking this discriminating component to choice confidence, we considered the Medium-confidence trials. Importantly, these trials can be regarded as “unseen” data, as they are independent from those used to train the classifier. We subjected these trials through the same neural generators (i.e. spatial projections) estimated during discrimination of High vs. Low confidence trials and, as expected from a graded quantity, found that the mean component amplitudes for Medium-confidence trials were situated between, and significantly different from, those in the High- and Low-confidence trial groups (both p < .001, Fig. 2B). To verify whether modulation of the discriminant component amplitude by confidence might be purely explained by subjects’ performance on the task (i.e., accuracy of the direction decision), we conducted the same comparison using only correct trials, and showed that this pattern persisted (both p < .001, Fig. 2C). This additional test was motivated by the observation that trial-to-trial discriminant component amplitudes were positively correlated with decision accuracy (subject-averaged R = .13; one-sample t-test, t(23) = 8.6, p < .001, Fig. 2D), in line with an increase in confidence with performance.

Further we addressed the possibility that the observed variability in the confidence discriminating component could be attributed to local fluctuations in attention, by conducting a serial autocorrelation analysis. As before, this model only explained a small fraction of the variance in component amplitudes (subject-averaged R^2^ = .03). We also assessed the influence of a neural signal known to correlate with attention (Thut et al., 2006) and predict visual discrimination (van Dijk et al., 2008), namely occipitoparietal prestimulus alpha power. To do this, we separated trials into High vs. Low alpha power groups, individually for each subject, and compared the corresponding average discriminant component amplitudes. We found that these did not differ significantly between the two groups (paired t-test, p=.19, Fig. 2E). Finally, we note that variability in the confidence discriminant component was also independent of stimulus difficulty, as this was held constant across all trials. We further supported this by showing that discriminant component amplitudes between the two identical-stimulus experimental blocks were not significantly correlated (mean R = .02; one-sample t-test, p = .39).

### fMRI correlates of reported confidence

To spatially characterise confidence signals in the fMRI data, we employed a general linear model approach (GLM). While this analysis was primarily aimed at identifying activation correlating with endogenous signatures of confidence derived from our EEG analysis at the time of the perceptual decision, our design matrix also included regressors accounting for variance linked to subjects’ behavioural confidence reports, as well as other potentially confounding factors (task performance, response time, attention, and visual stimulation; see Materials and methods).

**Figure 3.**
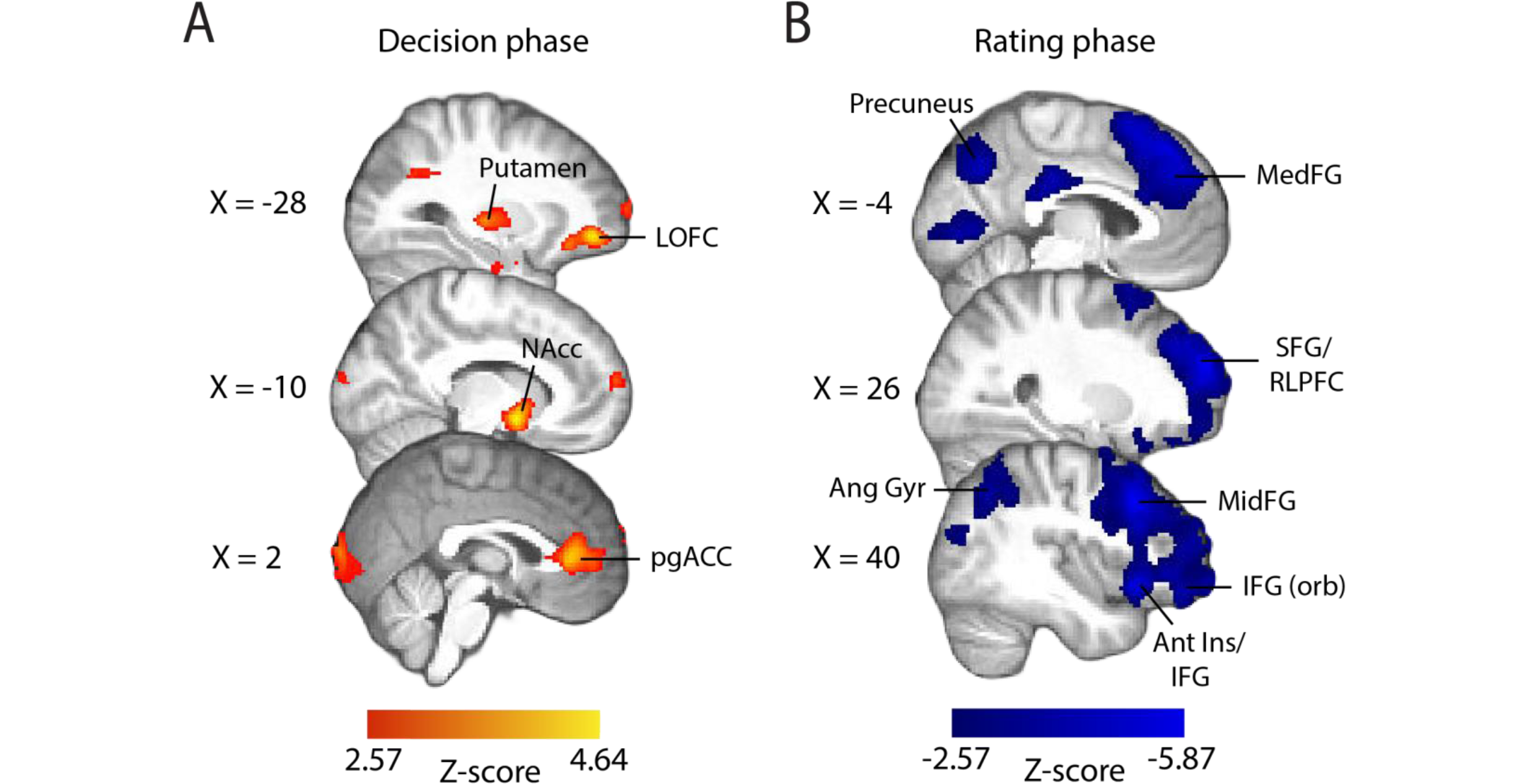
Parametric modulation of the BOLD signal by reported confidence. **A**, Clusters showing positive correlation with confidence during the decision phase of the trial. **B**, Clusters showing negative correlation with confidence at the onset of the rating cue (i.e., rating phase). All results are reported at |Z|≥2.57, and cluster-corrected using a resampling procedure (minimum cluster size 162 voxels; see Materials and Methods). *Ang Gyr,* angular gyrus; *Ant Ins,* anterior insula; *IFG (orb),* inferior frontal gyrus (orbital region); *LOFC,* lateral orbitofrontal cortex; *MedFG,* medial frontal gyrus; *MidFG,* middle frontal gyrus; *NAcc,* nucleus accumbens; *pgACC,* pregenual anterior cingulate cortex; *RLPFC,* rostrolateral prefrontal cortex; *SFG,* superior frontal gyrus. The complete lists of activations are shown in Tables 1 and 2.

Thus, we first inspected the activation patterns associated with confidence ratings during the perceptual decision phase of the trial (Fig. 3A). The coordinates of all activations are listed in Table 1. We found that the BOLD response increased with reported confidence in the striatum, lateral orbitofrontal cortex (OFC), the ventral anterior cingulate cortex (ACC) – areas thought to play a role in human valuation and reward (O’Doherty, 2004; Rushworth et al., 2007; Grabenhorst and Rolls, 2011) – as well as the right anterior middle frontal gyrus, amygdala/hippocampus, and visual association areas. Overall, these activations appear consistent with findings from previous studies that have identified spatial correlates of decision confidence (Rolls et al., 2010; De Martino et al., 2013; Heereman et al., 2015; Hebart et al., 2016). Negative activations (i.e., regions showing increasing BOLD response with decreasing reported confidence) were found in the right supplementary motor area, dorsomedial prefrontal cortex, right inferior frontal gyrus (IFG), anterior insula/frontal operculum, in line with previous reports of decision uncertainty near the time of decision (Heereman et al., 2015; Hebart et al., 2016).

**Table 1.** Complete list of brain activations correlating with subjects’ confidence reports, at the time of stimulus onset (decision phase).

| Brain region | BA | Laterality | Peak MNI coordinates (mm) |  |  | Z value (peak) |
| --- | --- | --- | --- | --- | --- | --- |
|  |  |  | X | Y | Z |  |
| Positive parametric effect (Z > 2.57) |  |  |  |  |  |  |
| Striatum (nucleus accumbens / ventral putamen) | - | L | -10 | 4 | -10 | 4.64 |
|  | - | R | 12 | 4 | -10 | 4.09 |
| Lateral orbitofrontal cortex | 11/47 | L | -28 | 46 | -8 | 4.46 |
|  | 47 | R | 32 | 38 | -6 | 3.86 |
| Anterior cingulate cortex | 32/10 | R, L | 2 | 36 | 6 | 4.19 |
| Lateral occipital cortex (inferior) | 19 | L | -42 | -68 | -10 | 4.04 |
|  | 19 | R | 48 | -82 | 8 | 3.13 |
| Middle frontal gyrus (anterior) | 10 | R | 40 | 62 | 10 | 3.94 |
| Striatum (dorsal putamen / pallidum) | - | L | -28 | -18 | 2 | 3.72 |
| Occipital pole | 17 | R, L | 4 | -102 | 8 | 3.66 |
| Cerebellum | - | R | 22 | -46 | -22 | 3.55 |
| Inferior temporal gyrus | 37 | R | 54 | -46 | -16 | 3.51 |
| Negative parametric effect (Z < -2.57) |  |  |  |  |  |  |
| Superior frontal gyrus (supplementary motor area) | 6 | R | 14 | 12 | 64 | 5.62 |
| Dorsomedial prefrontal cortex | 6/32 | R, L | -6 | 12 | 52 | 4.13 |
| Inferior frontal gyrus | 44/45 | R | 50 | 16 | 2 | 3.95 |
| Precentral gyrus | 6 | R | 50 | 4 | 46 | 3.61 |
|  | 6 | L | -44 | 2 | 38 | 3.46 |

MNI, Montreal Neurological Institute; L, left hemisphere; R, right hemisphere; BA, approximate Broadmann area.

During the metacognitive report stage of the trial (i.e., rating phase, Fig. 3B), we found negative correlations with confidence ratings in extended networks (Table 2) which included regions of the rostrolateral prefrontal cortex (bilateral, right lateralised), middle frontal gyrus, superior frontal gyrus (extending along the cortical midline and into the medial prefrontal cortex), orbital regions of the IFG, angular gyrus, precuneus, posterior cingulate cortex (PCC), and regions of the occipital and middle temporal cortices. These activations are largely in line with research on the spatial correlates of choice uncertainty (Grinband et al., 2006; Fleming et al., 2012) and metacognitive evaluation (Fleming et al., 2012; Molenberghs et al., 2016). Finally, positive correlations were observed in the striatum and amygdala/hippocampus, as well as motor cortices. Intriguingly, the seemingly distinct confidence-related network activations at the time of the perceptual decision vs. metacognitive report suggest these regions may encode qualitatively distinct representations of confidence at different times within the trial, for example faster and more automated representations of confidence (see (Lebreton et al., 2015)) around the time of decision, in contrast to metacognitive representations, when explicit evaluation/report are required.

**Table 2.** Complete list of brain activations correlating with subjects’ confidence reports, at the time of confidence rating (rating phase).

| Brain region | BA | Laterality | Peak MNI coordinates (mm) |  |  | Z value (peak) |
| --- | --- | --- | --- | --- | --- | --- |
|  |  |  | X | Y | Z |  |
| Positive parametric effect (Z > 2.57) |  |  |  |  |  |  |
| Amygdala / Hippocampus | - | R | 28 | -10 | -14 | 4.16 |
|  | - | L | -28 | -12 | -12 | 3.27 |
| Putamen | - | L | -22 | 18 | 2 | 4.01 |
| Precentral gyrus | 6/4 | L | -38 | -10 | 70 | 3.87 |
|  | 6 | R | 38 | -14 | 70 | 3.04 |
| Negative parametric effect (Z < -2.57) |  |  |  |  |  |  |
| Angular gyrus | 39 | L | -58 | -56 | 34 | 5.87 |
| Angular gyrus | 39 | R | 60 | -54 | 36 | 5.82 |
| Superior frontal gyrus / RLPFC | 10/9 | R | 24 | 58 | 26 | 5.84 |
|  | 10/9 | L | -20 | 52 | 26 | 5.2 |
| Inferior frontal gyrus (orbital area) / Anterior insula | 13/45 | L | -44 | 24 | -8 | 5.58 |
|  | 13/45 | R | 42 | 22 | -6 | 5.26 |
| Middle frontal gyrus | 8/9 | R | 44 | 20 | 42 | 5.56 |
|  | 8/9 | L | -40 | 20 | 42 | 4.92 |
| Medial frontal gyrus | 8/9 | L, R | 0 | 42 | 34 | 5.19 |
| Inferior frontal gyrus (triangular area) | 45 | L | -50 | 22 | 6 | 5.02 |
|  | 45 | R | 58 | 30 | 8 | 4.94 |
| Precuneus | 7 | L, R | -2 | -68 | 38 | 4.51 |
| Occipitotemporal gyrus | 37 | L | -38 | -62 | -22 | 4.34 |
| Posterior cingulate cortex | 23 | L, R | -2 | -26 | 32 | 4.76 |
| Middle temporal gyrus (anterior) | 20/21 | R | 50 | 2 | -34 | 4.36 |
| Thalamus | - | R | 10 | -10 | 2 | 4.35 |
|  | - | L | -12 | -10 | 6 | 3.82 |
| Lingual gyrus | 18 | L | -2 | -80 | 0 | 4.14 |
| Calcarine cortex | 17 | R | 16 | -90 | 2 | 4.14 |
|  | 17 | L | -12 | -92 | 2 | 3.93 |
| Middle temporal gyrus (posterior) | 21/37 | R | 56 | -34 | -12 | 3.93 |
|  | 21 | L | -54 | -30 | -8 | 3.82 |
| Inferior occipital gyrus | 18 | R | 28 | -90 | -10 | 3.19 |
| Lateral occipital cortex (superior) | 19 | R | 44 | -74 | 20 | 3.58 |
|  | 19 | L | -40 | -88 | 20 | 3.43 |

MNI, Montreal Neurological Institute; L, left hemisphere; R, right hemisphere; BA, approximate Broadmann area; RLPFC, rostrolateral prefrontal cortex

### fMRI correlates of EEG-derived confidence

We used the single-trial variability associated with the confidence discriminating component to construct a parametric EEG-derived fMRI regressor (Y_CONF_ regressor), in order to identify potential brain regions encoding internal representations of early confidence as captured by this EEG component.

Crucial to our approach was modelling the fMRI activation using time-resolved, electrophysiologically-derived signatures of confidence which were specific to each subject. These measures captured the variability in the neural representation of confidence around the perceptual decision itself (i.e., prior to behavioural response), and at a time point of maximum confidence discrimination, thus allowing us to detect its associated spatial correlates with increased temporal and spatial precision, relative to what behavioural ratings and fMRI measurements alone permitted. Importantly, as these signals were only partially correlated with reported confidence, they could potentially provide additional explanatory power in our fMRI model.

This EEG-informed fMRI analysis revealed a large cluster in the ventromedial prefrontal cortex (VMPFC, peak MNI coordinates [-8 40 -14]), extending into the subcallosal region and ventral striatum, and a smaller cluster in the right precentral gyrus (peak MNI coordinates [30 -20 64]), where the BOLD response correlated positively with the EEG-derived confidence discriminating component (Fig. 4). Recent studies have linked the VMPFC to confidence in value-based as well as other complex decisions (De Martino et al., 2013; Lebreton et al., 2015), however this region is not typically associated with confidence in perceptual decisions (though see (Heereman et al., 2015)). This finding is consistent with recent work proposing a domain-general role for the VMPFC in encoding confidence (Lebreton et al., 2015), and raises the possibility that this region holds information about early confidence signals emerging prior to the execution of a behavioural choice.

Importantly, we note that the EEG-derived measures, which informed the fMRI analysis, were independent of task difficulty, accuracy, or attention, as discussed in previous sections. Additionally, the GLM model included separate regressors controlling for these variables, and other potential confounds (see Materials and Methods). In particular, our simultaneous EEG/fMRI approach allowed the introduction of an additional level of control for attentional confounds in the fMRI analysis, namely by including the same EEG-derived index of attention as a nuisance predictor in the GLM model. This regressor showed significant correlation with the intraparietal regions and the frontal eye fields, consistent with the dorsal attentional network thought to be involved in top-down control of visual attention (Corbetta and Shulman, 2002).

**Figure 4.**
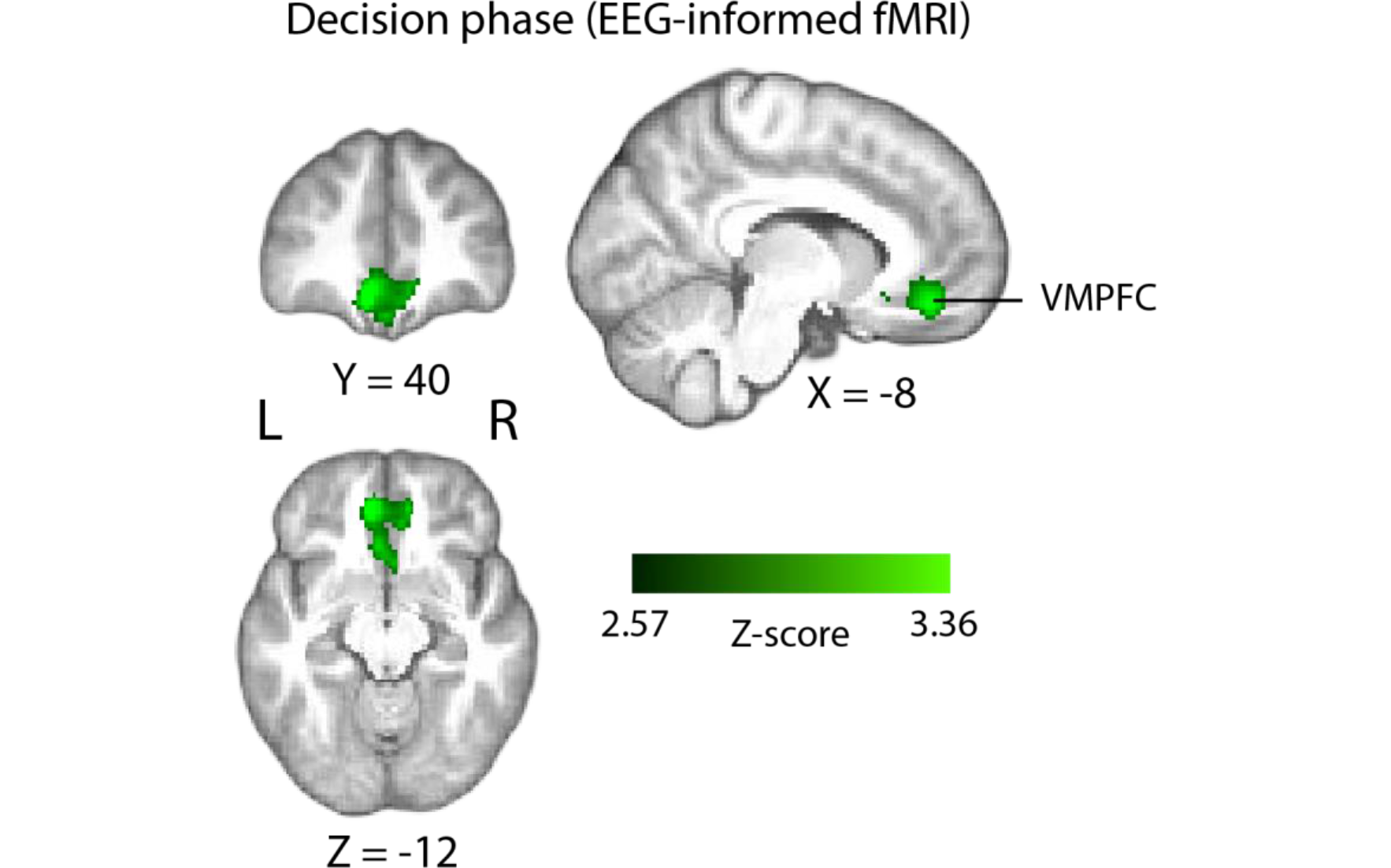
Positive parametric modulation of the BOLD signal by EEG-derived single-trial confidence measures (see Materials and Methods), during the decision phase of the trial. Results are reported at |Z|≥2.57, and cluster-corrected using a resampling procedure (minimum cluster size 162 voxels). *VMPFC,* ventromedial prefrontal cortex.

Next, we asked whether BOLD activation observed in the VMPFC during the perceptual decision period was uniquely associated with the EEG-derived Y_CONF_ regressor, i.e., over and above what could be explained by the behavioural confidence ratings (i.e., the Ratings_DEC_ regressor, Fig. 3A) alone. To test this, we compared mean parameter estimates (z-scored beta values) associated with the two predictors, within the VMPFC region identified with the Y_CONF_ regressor. We found that, across subjects, these were significantly higher for the Y_CONF_ regressor than for the Ratings_DEC_ regressor (paired t-test, t (23) = 9.48, p<.001). Moreover, VMPFC parameter estimates for the Y_CONF_ regressor remained significantly higher than those associated with the Ratings_DEC_, even when the latter were obtained with a control GLM model that did not include Y_CONF_ as a predictor (paired t-test, t (23) = 7.99, p<.001). Taken together, these observations indicate that our EEG-derived endogenous measures of confidence were better predictors of VMPFC activity at the time of decision than the post-decision behavioural reports.

Interestingly, the scalp map associated with our confidence discriminating EEG component showed a diffused topography including contributions from several centroparietal electrode sites. This emphasizes that spatial inferences made on the basis of scalp maps alone are limited due to volume conduction effects and field spread on the scalp-measured EEG signals (thus further highlighting the utility of the simultaneous EEG/fMRI method in locating these signals). This being considered, another possibility is that the observed spatial pattern reflects sources of shared variance between the EEG component and confidence ratings themselves (which was otherwise controlled for in our original fMRI analysis). To test this, we ran a separate control GLM analysis where the confidence ratings (Ratings_DEC_) regressor was removed, and found that with this model the Y_CONF_ regressor explained additional variability of the BOLD signal within several regions, including precuneus/PCC regions of the parietal cortex. Notably, these regions have been previously shown to scale with confidence (De Martino et al., 2013; White et al., 2014) and hypothesised to play a role in metacognitive ability (McCurdy et al., 2013).

### Psychophysiological interaction (PPI) analysis

Having identified the VMPFC as uniquely encoding a confidence signal early on in the trial (i.e., near the time of the perceptual decision), we next sought to examine potential functional interactions with regions holding neural representations of confidence at later stages, during explicit metacognitive evaluation/report. To this end, we conducted a functional connectivity analysis, using a psychophysiological interaction (PPI) approach (see Materials and methods). The PPI allows one to look for regions in the brain where the fMRI
response is modulated by the interaction between ongoing activation in a seed region (i.e., a physiological variable) and an experimental condition (i.e., psychological variable) (O’Reilly et al., 2012). Here, we used the VMPFC as a seed and looked for regions across the brain in where BOLD activation time series correlated with that of the seed during the perceptual decision phase of the trial (i.e., defined here as the trial-by-trial time window between onset of the motion stimulus and subject’s explicit commitment to choice). As we were primarily interested in interactions with networks involved in confidence processing during explicit metacognitive evaluation, we performed a conjunction analysis to examine the spatial overlap with these regions (as informed by our previous GLM analysis, see Fig. 3B). Note that during the rating phase of the trial, we observed predominantly negative correlations with confidence (whereas the EEG-derived confidence regressor at the time of choice revealed *positive* correlations with VMPFC signals). Accordingly, we sought to identify negative couplings with VMPFC activity in the PPI. Indeed, this analysis revealed two clusters in the left anterior prefrontal cortex (aPFC, peak MNI coordinates [-40 46 4]) and right dorsolateral prefrontal cortex (dlPFC, peak MNI coordinates [48 22 30]), respectively (Fig. 5), which showed increased negative correlation with VMPFC activation during the perceptual decision.

**Figure 5.**
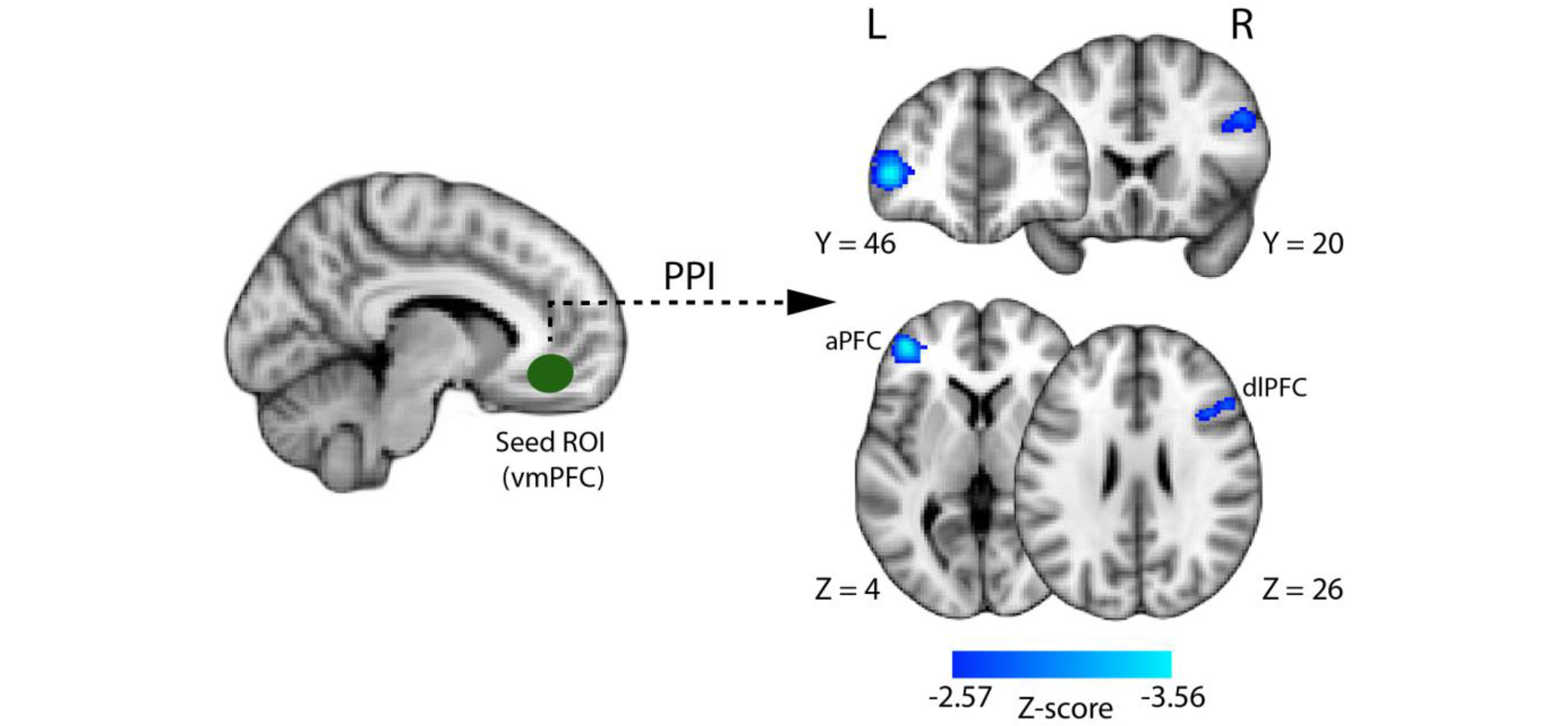
Psychophysiological interaction (PPI) analysis showing functional connectivity with the ventromedial prefrontal cortex (i.e., the seed region of interest; approximate location shown in green) during the perceptual decision . Displayed here are PPI activations that overlap spatially with those regions showing parametric modulation with subjective confidence at the time of metacognitive report (see Fig. 3B). Two clusters in the anterior and dorsolateral prefrontal cortices, respectively (shown in blue), show stronger negative correlation with the VMPFC during the perceptual decision. All results are reported at |Z|≥2.57, and cluster-corrected using a resampling procedure (minimum cluster size 162 voxels).

## Discussion

Here, we used a simultaneous EEG/fMRI approach to investigate the neural correlates of confidence during perceptual decisions. We found that BOLD activation in a region of the VMPFC was uniquely explained by the single-trial variability in an early EEG-derived neural signature of confidence occurring prior to subjects’ behavioural expression of response. Importantly, we showed that this activity was additional to what could be explained by subjects’ behavioural reports alone. Our results provide empirical support for the involvement of the VMPFC in confidence of perceptual decisions, consistent with recent evidence for a domain-general role of the VMPFC in encoding decision confidence. In turn this suggests that the VMPFC may support an early readout of confidence that precedes explicit metacognitive evaluation.

Our method allowed us to capitalise on the increased explanatory power inherent to our time-resolved internal measures of confidence, to identify relevant activation in the fMRI data. This, in turn, provided a more precise spatiotemporal characterisation than allowed by fMRI measures alone. The observation that the VMPFC holds information about confidence signals occurring prior to behavioural response is intriguing, as it raises novel possibilities for the role of this region in the confidence processing stream. Specifically, the VMPFC may encode early confidence representations (at, or near, the time of decision), which in turn could have important adaptive functions in influencing action that follows from the perceptual decision, and potentially informing choice itself (Lak et al., 2017). Additionally, such signals may be qualitatively different from confidence estimates available at the time of report as the latter are likely to undergo additional processing that continues after a choice is made (Resulaj et al., 2009; Pleskac and Busemeyer, 2010; Moran et al., 2015).

Interestingly, computational/neurobiological accounts of confidence processing have proposed architectures by which a first-level form of confidence in a decision emerges as a natural property of the same neural processes that support the decision, which in turn is read out (i.e., summarised) by separate higher-order monitoring network(s) (Insabato et al., 2010; Meyniel et al., 2015; Pouget et al., 2016). As the VMPFC is not typically known to support perceptual decision processes, the VMPFC confidence signals we observe here are thus likely to represent a readout of confidence-related information.

Consistent with a role as a monitoring module providing a confidence readout, recent work suggests the VMPFC may encode confidence in a task-independent and possibly domain-general manner. Specifically, several functional neuroimaging studies have shown positive modulation of VMPFC activation by confidence, across a range of decision making tasks (Rolls et al., 2010; De Martino et al., 2013; Heereman et al., 2015; Lebreton et al., 2015). Notably, one study showed that fMRI activation in the VMPFC was modulated by confidence across four different tasks involving both value-based and non-value based rating judgments (Lebreton et al., 2015). Furthermore, evidence from memory-related decision making research appears to also implicate the VMPFC in confidence processing (see Hebscher and Gilboa (2016) for a review). Our work, therefore, complements the existing literature by bringing empirical support for the involvement of VMPFC in perceptual decision making.

Nonetheless, confidence representations in the VMPFC may in themselves not be sufficient or available in an appropriate form for metacognitive report (Grimaldi et al., 2015), thus requiring further read-out from higher-order networks (Insabato et al., 2010) such as the prefrontal cortex (Fleming et al., 2012; De Martino et al., 2013). In line with this hypothesis, we showed evidence for a functional interaction between the VMPFC and regions of the prefrontal cortex that hold neural representations of confidence during explicit metacognitive reports. This functional link was observed in the perceptual decision phase of the trial, potentially indicative of an early transfer of confidence-related information to higher-order regions for metacognitive appraisal.

The observation that the VMPFC, a region known for its involvement in choice-related subjective valuation (Philiastides et al., 2010; Rangel and Hare, 2010; Bartra et al., 2013; Pisauro et al., 2017) encodes confidence signals during perceptual decisions raises an interesting possibility for interpreting our results. Our behavioural paradigm did not involve an explicit reward/feedback manipulation and accordingly, the observed confidence-related activation cannot be interpreted as an externally driven value signal. Instead, as has been suggested previously (Barron et al., 2015; Lebreton et al., 2015), a likely explanation is that, by being an internal measure of performance accuracy, confidence is inherently valuable. Such a signal may represent *implicit* reward and possibly act as a teaching signal (Daniel and Pollmann, 2012; Guggenmos et al., 2016; Lak et al., 2017) to drive learning (e.g., perceptual learning (Law and Gold, 2009; Kahnt et al., 2011; Diaz et al., 2017).

In line with this interpretation, Hebart et al. (2016) observed positive correlation with confidence in the ventral striatum, a region known for its involvement in reward (O’Doherty et al., 2004). Authorssuggest that confidence signals in this region may play a role in confidence-driven learning, such that feelings of reward associated with a choice reinforce optimal behavior on subsequent choices. A different study (Guggenmos et al., 2016) demonstrated that regions of the human mesolimbic dopamine system, namely the striatum and ventral tegmental area, encoded both anticipation and prediction error related to decision confidence (i.e., in the absence of feedback), similar to what is typically observed during reinforcement learning tasks where feedback is explicit (Preuschoff et al., 2006; Fouragnan et al., 2015; Fouragnan et al., In Press). Importantly, these effects were predictive of subjects’ perceptual learning efficiency. Thus, confidence in valuation/reward networks could be propagated back to the decision systems to optimize the dynamics of the decision process, possibly by means of a reinforcement-learning mechanism.

In conclusion, we showed that by employing a simultaneous EEG/fMRI approach, we were able to localise an early representation of confidence in the brain with higher spatiotemporal precision than allowed by fMRI alone. In doing so, we provided novel empirical evidence for the encoding of a generalised confidence readout signal in the VMPFC preceding explicit metacognitive report. Our findings provide a starting point for further investigations into the neural dynamics of confidence formation in the human brain and its interaction with other cognitive processes such as learning, and the decision itself.

## Materials and methods

### Participants

Thirty subjects participated in the simultaneous EEG/fMRI experiment. Four were subsequently removed from the analysis due to near chance (n=3) and ceiling (n=1) performance, respectively, on the perceptual discrimination task. Additionally, one subject was excluded whose confidence reports covered only a limited fraction of the provided rating scale, thus yielding an insufficient number of trials to be used in the EEG discrimination analysis (see below). Finally, one subject had to be removed due to poor (chance) performance of the EEG decoder (see below). All results presented here are based on the remaining 24 subjects (age range 20-32 years). All were right-handed, had normal or corrected to normal vision, and reported no history of neurological problems. The study was approved by the College of Science and Engineering Ethics Committee at the University of Glasgow (CSE01355) and informed consent was obtained from all participants.

### Stimuli and task

All stimuli were created and presented using the PsychoPy software (Peirce, 2007). They were displayed via an LCD projector (frame rate=60Hz) on a screen placed at the rear opening of the bore of the MRI scanner, and viewed through a mirror mounted on the head coil (distance to screen = 95cm). Stimuli consisted of random dot kinematograms (Newsome and Pare, 1988), whereby a proportion of the dots moved coherently to one direction (left vs. right), while the remainder of the dots moved at random. Specifically, each stimulus consisted of a dynamic field of white dots (number of dots=150; dot diameter=0.1 degrees of visual angle, dva; dot life time=4 frames; dot speed=6 dva/s), displayed centrally on a grey background through a circular aperture (diameter=6 dva). Task difficulty was controlled by manipulating the proportion of dots moving coherently in the same direction (i.e., motion coherence).

We aimed to maintain overall performance on the main perceptual decision task consistent across subjects (i.e., near perceptual threshold, at approximately 75% correct). For this reason, task difficulty was calibrated individually for each subject on the basis of a separate training session, prior to the day of the main experiment.

#### Training

To first familiarise subjects with the random dot stimuli and facilitate learning on the motion discrimination task, subjects first performed a short simplified version of the main task (lasting approx. 10 minutes), where feedback was provided on each trial. The task, which required making speeded direction discriminations of random dot stimuli (see below), began at a low-difficulty level (motion coherence = 40%) and gradually increased in difficulty in accordance with subjects’ online behavioural performance (a 3-down-1-up staircase procedure, where three consecutive correct responses resulted in a 5% decrease in motion coherence, whereas one incorrect response yielded a 5% increase). This was followed by a second, similar task, which served to determine subject-specific psychophysical thresholds. Seven motion coherence levels (5%, 8%, 12%, 18%, 28%, 44%, 70%) were equally and randomly distributed across 350 trials. The proportion of correct responses was separately computed for each motion coherence level, and a logarithmic function was fitted through the resulting values in order to estimate an optimal motion coherence yielding a mean performance of approximately 75% correct. Subjects who showed near-chance performance across all coherence levels or showed no improvement in performance with increasing motion coherence were not tested further and did not participate in the main experiment. No feedback was given for this or any of the subsequent tasks.

#### Main task

On the day of the main experiment, subjects practiced the main task once outside the scanner, and again inside the scanner prior to the start of the scan (a short 80 trial block each time). Subjects made left vs. right direction discriminations of random dot kinematograms and rate how confident they were in their choices, on a trial-by-trial basis (Fig. 1A).

Each trial began with a random dot stimulus lasting for a maximum of 1.2 s, or until the subject made a behavioural response. Subjects were instructed to respond as quickly as possible, and had a time limit of 1.35 s to do so. The message “Oops! Too slow” was displayed if this time limit was exceeded or no direction response was made. Once the dot stimulus disappeared, the screen remained blank until the 1.2 s stimulation period elapsed and through an additional random delay (1.5-4 s). Next, subjects were presented with a rating scale for 3 s, during which they reported their confidence in the previous direction decision. The confidence scale was represented intuitively by means of a white horizontal bar of linearly varying thickness, with the thick end representing high confidence. Its orientation on the horizontal axis (thin-to-thick vs. thick-to-thin) informed subjects of the response mapping, and this was equally and randomly distributed across trials to control for motor preparation effects. To make a confidence response, subjects moved an indicator (a small white triangle) along a 9-point marked line. The indicator changed colour from white to yellow when a confidence response was selected and this remained on the screen until the 3 s elapsed). A final delay (blank screen, jittered between 1.5-4 s) ended the trial. Failing to provide either a direction or a confidence response within the respective allocated time limits on a given trial rendered it invalid, and this was subsequently removed from further analyses. This resulted in a total fraction of .04 (.02 and .02, respectively) of trials being discarded.

Subjects performed 2 experimental blocks of 160 trials each, corresponding to two separate fMRI runs. Each block contained two short (30 s) rest breaks, during which the MR scanner continued to run. Subjects were instructed to remain still throughout the entire duration of the experiment, including during rest breaks and in between scans. Motion coherence was held constant across trials, at the subject-specific level estimated during training. The direction of the dots was equally and randomly distributed across trials. To control for confounding effects of low-level trial-to-trial variability in stimulus properties on decision confidence, an identical set of stimuli was used in the two experimental blocks. Specifically, for each subject, the random seed, which controlled dot stimulus motion parameters in the stimulus presentation software was set to a fixed value. This manipulation allowed for subsequent control comparisons between pairs of identical stimuli.

Subjects were encouraged to explore the entire scale when making their responses and to abstain from making a confidence response on a given trial if they became aware of having made a motor mapping error (this was in an effort to avoid “confident errors”). They were instructed to make their responses as quickly and accurately as possible, and provide a response on every trial. All behavioural responses were executed using the right hand, on an MR-compatible button box.

### EEG data acquisition

EEG data was collected using an MR-compatible EEG amplifier system (Brain Products, Germany). Continuous EEG data was recorded using the Brain Vision Recorder software (Brain Products, Germany) at a sampling rate of 5000 Hz. We used 64 Ag/AgCl scalp electrodes positioned according to the 10-20 system, and one nasion electrode. Reference and ground electrodes were embedded in the EEG cap and were located along the midline, between electrodes Fpz and Fz, and between electrodes Pz and Oz, respectively. Each electrode had in-line 10 kOhm surface-mount resistors to ensure subject safety. Input impedance was adjusted to <25 kOhm for all electrodes. Acquisition of the EEG data was synchronized with the MR data acquisition (Syncbox, Brain Products, Germany), and MR-scanner triggers were collected separately to enable offline removal of MR gradient artifacts from the EEG signal. Scanner trigger pulses were lengthened to 50μs using a built-in pulse stretcher, to facilitate accurate capture by the recording software. Experimental event markers (including participants’ responses) were synchronized, and recorded simultaneously, with the EEG data.

### EEG data processing

Preprocessing of the EEG signals was performed using Matlab (Mathworks, Natick, MA). EEG signals recorded inside an MR scanner are contaminated with gradient artifacts and ballistocardiogram (BCG) artifacts due to magnetic induction on the EEG leads. To correct for gradient-related artifacts, we constructed average artifact templates from sets of 80 consecutive functional volumes centred on each volume of interest, and subtracted these from the EEG signal. This process was repeated for each functional volume in our dataset. Additionally, a 12 ms median filter was applied in order to remove any residual spike artifacts. Further, we corrected for standard EEG artifacts and applied a 0.5–40 Hz band-pass filter in order to remove slow DC drifts and high frequency noise. All data were downsampled to 1000 Hz.

To remove eye movement artifacts, subjects performed an eye movement calibration task prior to the main experiment (with the MRI scanner turned off, to avoid gradient artifacts), during which they were instructed to blink repeatedly several times while a central fixation cross was displayed in the centre of the computer screen, and to make lateral and vertical saccades according to the position of the fixation cross. We recorded the timing of these visual cues and used principal component analysis to identify linear components associated with blinks and saccades, which were subsequently removed from the EEG data (Parra et al., 2005).

Next, we corrected for cardiac-related (i.e., ballistocardiogram, BCG) artifacts. As these share frequency content with the EEG, they are more challenging to remove. To minimise loss of signal power in the underlying EEG signal, we adopted a conservative approach by only removing a small number of subject-specific BCG components, using principal component analysis. We relied on the single-trial classifiers to identify discriminating components that are likely to be orthogonal to the BCG. BCG principal components were extracted from the data after the data were first low-pass filtered at 4 Hz to extract the signal within the frequency range where BCG artifacts are observed. Subject-specific principal components were then determined (average number of components across subjects: 1.8). The sensor weightings corresponding to those components were projected onto the broadband data and subtracted out. Finally, data were baseline corrected by removing the average signal during the 100 ms prestimulus interval.

### Single-trial EEG analysis

To increase statistical power of the EEG data analysis, trials were separated into three confidence groups (Low, Medium, High), on the basis of the original 9-point confidence rating scale. Specifically, we isolated High- and Low-confidence trials by pooling across each subject’s three highest and three lowest ratings, respectively. To ensure robustness of our single trial EEG analysis, we imposed a minimum limit of 50 trials per confidence trial group. For those data sets where subjects had an insufficient number of trials in the extreme ends of the confidence scale, neighbouring confidence bins were included to meet this limit.

We used a single-trial multivariate discriminant analysis, combined with a sliding window approach (Parra et al., 2005; Sajda et al., 2009) to discriminate between High and Low confidence trials in the stimulus-locked EEG data. This method aims to estimate, for predefined time windows of interest, an optimal combination of EEG sensor linear weights (i.e., a spatial filter) which, applied to the multichannel EEG data, yields a one-dimensional projection (i.e., a “discriminant component”) that maximally discriminates between the two conditions of interest. Importantly, unlike univariate trial-average approaches for event-related potential analysis, this method spatially integrates information across the multidimensional sensor space, thus increasing signal-to-noise ratio whilst simultaneously preserving the trial-by-trial variability in the signal, which may contain task-relevant information. In our data, we identified confidence-related discriminating components, ***y***(t), by applying a spatial weighting vector ***w*** to our multidimensional EEG data ***x***(t), as follows:

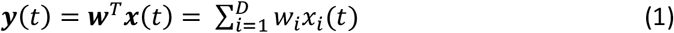

where *D* represents the number of channels, indexed by, and indicates the transpose of the matrix. To estimate the optimal discriminating spatial weighting vector ***w***, we used logistic regression and a reweighted least squares algorithm (Jordan and Jacobs, 1994). We applied this method to identify ***w*** for short (60 ms) overlapping time windows centred at 10 ms-interval time points, between -100 and 1000 ms relative to the onset of the random dot stimulus (i.e., the perceptual decision phase of the trial). This procedure was repeated for each subject and time window. Applied to an individual trial, spatial filters (***w***) obtained this way produce a measurement of the discriminant component amplitude for that trial. In separating the High and Low trial groups, the discriminator was designed to map the component amplitudes for one condition to positive values and those of the other condition to negative values. Here, we mapped the High confidence trials to positive values and the Low confidence trials to negative values, however note that this mapping is arbitrary.

To quantify the performance of the discriminator for each time window, we computed the area under a receiver operating characteristic (ROC) curve (i.e., the Az value), using a leave-one-out trial procedure (Duda et al., 2001). We determined significance thresholds for the discriminator performance using a bootstrap analysis whereby trial labels were randomised and submitted to a leave-one-out test. This randomisation procedure was repeated 500 times, producing a probability distribution for *Az,* which we used as reference to estimate the *Az* value leading to a significance level of p<0.01.

Given the linearity of our model we also computed scalp projections of the discriminating components resulting from Eq. 1 by estimating a forward model for each component:

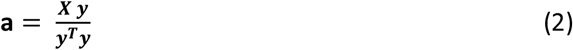

where the EEG data (***X***) and discriminating components (***y***) are now in a matrix and vector notation, respectively, for convenience (i.e., both ***X*** and ***y*** now contain a time dimension). Equation 2 describes the electrical coupling of the discriminating component that explains most of the activity in ***X***. Strong coupling indicates low attenuation of the component and can be visualised as the intensity of vector **a**.

### Single-trial power analysis

We calculated prestimulus alpha power (8-12Hz) in the 400 ms epoch beginning at -500 ms relative to the onset of the random dot stimulus. To do this, we used the multitaper method (Mitra and Pesaran, 1999) as implemented in the FieldTrip toolbox for Matlab (http://www.ru.nl/neuroimaging/fieldtrip). Specifically, for each epoch data were tapered using discrete prolate spheroidal sequences (2 tapers for each epoch; frequency smoothing of ±4Hz) and Fourier transformed. Resulting frequency representations were averaged across tapers and frequencies. Single-trial power estimates were then extracted from the occipitoparietal sensor with the highest overall alpha power and baseline normalised through conversion to decibel units (dB).

### MRI data acquisition

Imaging was performed at the Centre for Cognitive Neuroimaging, Glasgow, using a 3-Tesla Siemens TIM Trio MRI scanner (Siemens, Erlangen, Germany) with a 12-channel head coil. Cushions were placed around the head to minimize head motion. We recorded two experimental runs of 794 whole-brain volumes each, corresponding to the two blocks of trials in the main experimental task. Functional volumes were acquired using a T2^*^-weighted gradient echo, echo-planar imaging sequence (32 interleaved slices, gap: 0.3 mm, voxel size: 3 × 3 × 3 mm, matrix size: 70 × 70, FOV: 210 mm, TE: 30 ms, TR: 2000 ms, flip angle: 80°). Additionally, a high-resolution anatomical volume was acquired at the end of the experimental session using a T1-weighted sequence (192 slices, gap: 0.5 mm, voxel size: 1 × 1 × 1 mm, matrix size: 256 × 256, FOV: 256 mm, TE: 2300 ms, TR: 2.96 ms, flip angle: 9°), which served as anatomical reference for the functional scans.

### fMRI preprocessing

The first 10 volumes prior to task onset were discarded from each fMRI run to ensure a steady-state MR signal. Additionally, 13 volumes were discarded from the post-task period at the end of each block. The remaining 771 volumes were used for statistical analyses. Pre-processing of the MRI data was performed using the FEAT tool of the FSL software (http://www.fmrib.ox.ac.uk/fsl) and included slice-timing correction, high-pass filtering (>100 s), and spatial smoothing (with a Gaussian kernel of 8 mm full width at half maximum), and head motion correction (using the MCFLIRT tool). The motion correction preprocessing step generated motion parameters which were subsequently included as regressors of no interest in the general linear model (GLM) analysis (see fMRI analysis below). Brain extraction of the structural and functional images was performed using the Brain Extraction tool (BET). Registration of EPI images to standard space (Montreal Neurological Institute, MNI) was performed using the Non-linear Image Registration Tool with a 10-mm warp resolution. The registration procedure involved transforming the EPI images into an individual’s high-resolution space (with a linear, boundary-based registration algorithm, (Greve and Fischl, 2009)) prior to transforming to standard space. Registration outcome was visually checked for each subject to ensure correct alignment.

### fMRI analysis

Whole-brain statistical analyses of functional data were conducted using a general linear model (GLM) approach, as implemented in FSL (FEAT tool):

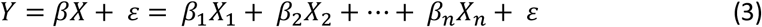

where *Y* represents the BOLD response time series for a given voxel, structured as a T×1 (T time samples) column vector, and represents the T×N (N regressors) design matrix, with each column representing one of the psychological regressors (see GLM analysis below for details), convolved with a canonical hemodynamic response function (double-gamma function). *β* represents the parameter estimates (i.e., regressor betas) resulting from the GLM analysis in the form of a N × 1 column vector. Lastly, ε is a T × 1 column vector of residual error terms. A first-level analysis was performed to analyse each subject’s individual runs. These were then combined at the subject-level using a second-level analysis (fixed effects). Finally, a third-level mixed-effects model (FLAME 1) was used to combine data across all subjects.

### Simultaneous EEG/fMRI analysis

With the combined EEG/fMRI approach, we sought to identify confidence-related activation in the fMRI surpassing what could be explained by the relevant behavioural predictors alone. In particular, we looked for brain regions where BOLD responses correlated with the confidence-discriminating component derived from the EEG analysis. Our primary motivation behind this approach was the hypothesis that endogenous trial-by-trial variability in the confidence discriminating EEG component (near the time of perceptual decision, and prior to behavioural response) would be more reflective of early internal representations of confidence at the single-trial level, compared to the metacognitive reports which are provided post-decisionally and therefore likely to be subjected to additional processes. We predicted that the simultaneous EEG/fMRI approach would enable identification of latent brain states that might remain unobserved with a conventional analysis approach. To this end, we extracted trial-by-trial amplitudes of ***y***(*t*) (resulting from Eq. 1) at the time window of maximum confidence discrimination, and used these to build a BOLD predictor, which we henceforth refer to as the Y_CONF_ regressor. Importantly, to avoid possible confounding effects of motor preparation/response, the time of this component was determined on a subject-specific basis, by only considering the period prior to the behavioural choice (mean peak discrimination time = 708 ms from stimulus onset, SD=162 ms). Thus, on average this was selected 287ms (SD=171 ms) prior to each subject’s mean response time.

Note that the trial-by-trial variability in our EEG component amplitudes is driven mostly by cortical regions found in close proximity to the recording sensors and to a lesser extent by distant (e.g., subcortical) structures. Nonetheless, an advantage of our EEG-informed fMRI predictors is that they can also reveal relevant fMRI activations within deeper structures, provided that their BOLD activity covaries with that of the cortical sources of our EEG signal.

### GLM analysis

We designed our GLM model to account for variance in the BOLD signal at two key stages of the trial, namely the perceptual decision period (beginning at the onset of the random dot visual stimulus) and the metacognitive evaluation/rating (beginning at the onset of the rating scale display), respectively. A total of 10 regressors were included in the model. Our primary predictor of interest was the EEG-derived endogenous measure of confidence (Y_CONF_ regressor). We modelled this as a stick function (duration = 0.1 s) locked to the stimulus onset, with event amplitudes parametrically modulated by the trial-to-trial variability in the confidence discriminating component ***y***(t). To ensure variance explained by this regressor was unique (i.e., not explained by subjects’ behavioural reports), we included a second regressor whose event amplitudes were parametrically modulated by confidence ratings, and which was otherwise identical to the Y_CONF_ regressor (i.e., Ratings_DEC_ regressor, duration = 0.1 s, locked to stimulus onset). Importantly, Y_CONF_ amplitudes were only moderately correlated with behavioural confidence ratings (mean R=.39, SD=.07), thus allowing us to exploit additional explanatory power inherent to this regressor. Other regressors of no interest for the perceptual decision stage included: one regressor parametrically modulated by prestimulus alpha power in the EEG signal (to control for potential attentional baseline effects), one categorical regressor (1/0) accounting for variability in response accuracy, and one unmodulated regressor (all event amplitudes set to 1) modelling stimulus-related visual responses of no interest across both valid and non-valid (missed) trials (all event durations = 0.1 s, locked to stimulus onset). To control for motor preparation/response, we also included a parametric regressor modulated by subjects’ reaction time on the direction discrimination task (duration = 0.1 s, locked to the time of behavioural response).

Additionally, locked to the onset of the metacognitive rating period, we included one parametric regressor (duration = 0.1 s) with event amplitudes modulated by subjects’ confidence ratings, one boxcar regressor with duration equivalent to subjects’ active behavioural engagement in confidence rating (to minimise effects relating to motor processes), and one unmodulated regressor (duration = 0.1 s). Lastly, we included one categorical boxcar regressor (1/0) to model non-task activation (i.e., rest breaks within each run). Motion correction parameters obtained from fMRI preprocessing were entered as additional covariates of no interest.

### Resampling procedure for fMRI thresholding

To estimate a significance threshold for our fMRI statistical maps whilst correcting for multiple comparisons, we performed a nonparametric permutation analysis that took into account the a priori statistics of the trial-to-trial variability in our primary regressor of interest (Y_CONF_), in a way that trades off cluster size and maximum voxel Z-score (Debettencourt et al., 2011). For each resampled iteration, we maintained the onset and duration of the regressor identical, whilst shuffling amplitude values across trials, runs and subjects. Thus, the resulting regressors for each subject were different as they were constructed from a random sequence of regressor amplitude events. This procedure was repeated 200 times. For each of the 200 resampled iterations, we performed a full 3-level analysis (run, subject, and group). Our design matrix included the same regressors of non-interest used in all our GLM analysis. This allowed us to construct the null hypothesis H_0_, and establish a threshold on cluster size and Z-score based on the cluster outputs from the permuted parametric regressors. Specifically, we extracted cluster sizes from all activations exceeding a minimal cluster size (5 voxels) and Z-score (2.57 per voxel) for positive correlations with the permuted parametric regressors. Finally, we examined the distribution of cluster sizes (number of voxels) for the permuted data and found that the largest 5% of cluster sizes exceeded 162 voxels. We therefore used these results to derive a corrected threshold for our statistical maps, which we then applied to the clusters observed in the original data (that is, Z=2.57, minimum cluster size of 162 voxels, corrected at p=0.05).

### Psychophysiological interaction analysis

We conducted a psychophysiological (PPI) analysis to explore potential functional connectivity between the region of the VMPFC found to uniquely explain trial-to-trial variability in our electrophysiologically-derived measures of confidence, and the rest of the brain, during the perceptual decision phase of the trial. To carry out the PPI analysis, we first extracted the time-series data from the seed region. Specifically, we identified the cluster of interest at the group level (i.e., in standard space) by applying the cluster correction procedure described in the previous section. Using this as a template, we constructed subject-specific masks of the voxels exhibiting the strongest correlation with the VMPFC region of interest, and back-projected these into the functional space of each individual. Resulting masks were used to compute average time-series data, separately for each subject and functional run, which subsequently served as the physiological regressor(s) in the PPI model. To carry out the PPI analysis, we performed a new GLM analysis. This included the following regressors, locked to the time of stimulus onset: (1) an unmodulated regressor (all event amplitudes set to 1), (2) the physiological regressor (time course of the VMPFC seed), (3) the psychological regressor (a boxcar function with event amplitudes set to 1 and duration parametrically modulated by subjects’ response time on the direction discrimination task), and (4) the interaction regressor. Additionally, motion parameters estimated during registration (see preprocessing step) were included as regressors of no interest. The statistical output from the interaction regressor thus reveals regions of the brain where correlation with the BOLD signal in the vmPFC is stronger during the perceptual decision than the rest of the trial. Importantly, this represents variance additional to that explained by the psychological and physiological regressors alone. Correction for multiple comparisons was performed on the whole brain using the outcome of the resampling procedure as described earlier. To identify only those activations which were common (i.e., overlapped spatially) with the confidence-related activations at the time of subjective reports (Fig. 3B), we created an intersection of the statistical maps associated with the PPI regressor and the Ratings_RAT_ regressor from the original GLM analysis. We only report resulting joint activations.

## Acknowledgements

This work was supported by the Economic and Social Research Council (ESRC; grant ES/L012995/1 to M.G.P.) and the British Academy (BA; grant SG121587 to M.G.P).

